# Assessing transcriptomic re-identification risks using discriminative sequence models

**DOI:** 10.1101/2023.04.13.536784

**Authors:** Shuvom Sadhuka, Daniel Fridman, Bonnie Berger, Hyunghoon Cho

## Abstract

Gene expression data provides molecular insights into the functional impact of genetic variation, for example through expression quantitative trait loci (eQTL). With an improving understanding of the association between genotypes and gene expression comes a greater concern that gene expression profiles could be matched to genotype profiles of the same individuals in another dataset, known as a linking attack. Prior works demonstrating such a risk could analyze only a fraction of eQTLs that are independent due to restrictive model assumptions, leaving the full extent of this risk incompletely understood. To address this challenge, we introduce the discriminative sequence model (DSM), a novel probabilistic framework for predicting a sequence of genotypes based on gene expression data. By modeling the joint distribution over all known eQTLs in a genomic region, DSM improves the power of linking attacks with necessary calibration for linkage disequilibrium and redundant predictive signals. We demonstrate greater linking accuracy of DSM compared to existing approaches across a range of attack scenarios and datasets including up to 22K individuals, suggesting that DSM helps uncover a substantial additional risk overlooked by previous studies. Our work provides a unified framework for assessing the privacy risks of sharing diverse omics datasets beyond transcriptomics.

## Introduction

The growing availability of large-scale genomic data repositories has led to increasing concerns for privacy of individuals from whom the data were collected [Bonomi et al., 2020, Naveed et al., 2015, Erlich and Narayanan, 2014]. While many nations and organizations have introduced policies and regulations (e.g., HIPAA and GDPR) to safeguard the collection, use, and sharing of personally-identifying information, existing policies often fall short of providing clear guidance regarding many widely-used types of biomedical data, including genetic sequences and functional genomic data, for which the underlying privacy risks are often unclear and only beginning to be understood [Wan et al., 2022, Clayton et al., 2019, Gürsoy et al., 2022b, Gürsoy et al., 2020, Bowler et al., 2022, Gürsoy et al., 2021]. This lack of guidance, particularly for functional genomic data, presents a key challenge for ensuring protection of study participants, leaving the possibility for future privacy breaches that may diminish public trust in the scientific community.

Transcriptomic data, such as gene expression measurements broadly shared in databases such as NCBI GEO [Barrett et al., 2010] and ArrayExpress [Athar et al., 2019], is a prominent example of biomedical data with incompletely understood privacy implications. While prior works [Schadt et al., 2012, Harmanci and Gerstein, 2016] have shown that the knowledge of *expression quantitative trait loci* (eQTLs)—i.e., genetic variants correlated with expression levels of a gene (referred to as eGenes)—could be used to extract genotypic information from gene expression profiles, the full extent of such leakage largely remains unknown. A key concern is that a malicious actor may exploit this leakage to carry out a *linking attack*, where one links an individual’s gene expression profile to their genotype profile in another dataset (or vice versa), potentially re-identifying the individual’s data and subsequently associating it with a sensitive attribute such as disease status.

Linking attacks require some criterion, which we call a match score function, to score and rank possible candidate matches between the gene expression and genotype datasets. The core challenge in accurately assessing the risk of a linking attack is therefore in devising the best possible match score function that could be leveraged by an attacker. Existing proposals for this function suffer from the key limitation that they account for only a small subset of (nearly) independent eQTLs as required by the simplified probabilistic models used in those approaches [Schadt et al., 2012, Harmanci and Gerstein, 2016]. As we will show, such restrictions have thus far obscured the extent of genotypic information leakage in expression data.

Here, we introduce the *discriminative sequence model* (DSM), which jointly models all known eQTLs in a genomic region to enable *sequence-level* inference of genotypes given a gene expression profile. DSM provides more accurate probability estimates for each candidate genotype profile originating from the same individual as the expression profile by calibrating for both correlation among genetic variants and redundant eQTL association signals. Using these probabilities as match scores, DSM leads to substantially greater success in simulated linking attacks than existing strategies in a wide range of scenarios including: (i) linking an expression profile against a large-scale candidate genotype set (e.g., including 22K individuals) representing a population whose ancestry background differs from the training set; (ii) linking when the membership of the target individual in the genotype set is unknown; and (iii) linking in the reverse direction—i.e., from a genotype profile to a panel of candidate expression profiles. Our work provides an essential tool for gaining a deeper understanding of privacy risks of transcriptomic data and its linkage with protected genetic information.

## Results

### Overview of the Discriminative Sequence Model (DSM)

We developed DSM based on the framework of conditional random fields [Sutton et al., 2012] to model the conditional distribution over the genotype sequence given a query gene expression profile. DSM facilitates linking of samples between gene expression and genotype datasets by scoring the likelihood of a target genetic sequence belonging to the same individual as the query gene expression profile (Fig. 1A,B). DSM extends the widely-used Li-Stephens hidden Markov model (HMM) of genetic sequences [Li and Stephens, 2003] to include additional probabilistic factors that capture the correlation between each eQTL and its corresponding set of known eGenes (genes whose expression is correlated with an eQTL). In contrast to existing models, DSM obtains calibrated probabilities accounting for all known eQTLs and their genotypic correlations (Fig. 1C,D). We detail our problem setting, threat model, and existing approaches in Methods.

**Figure 1:**
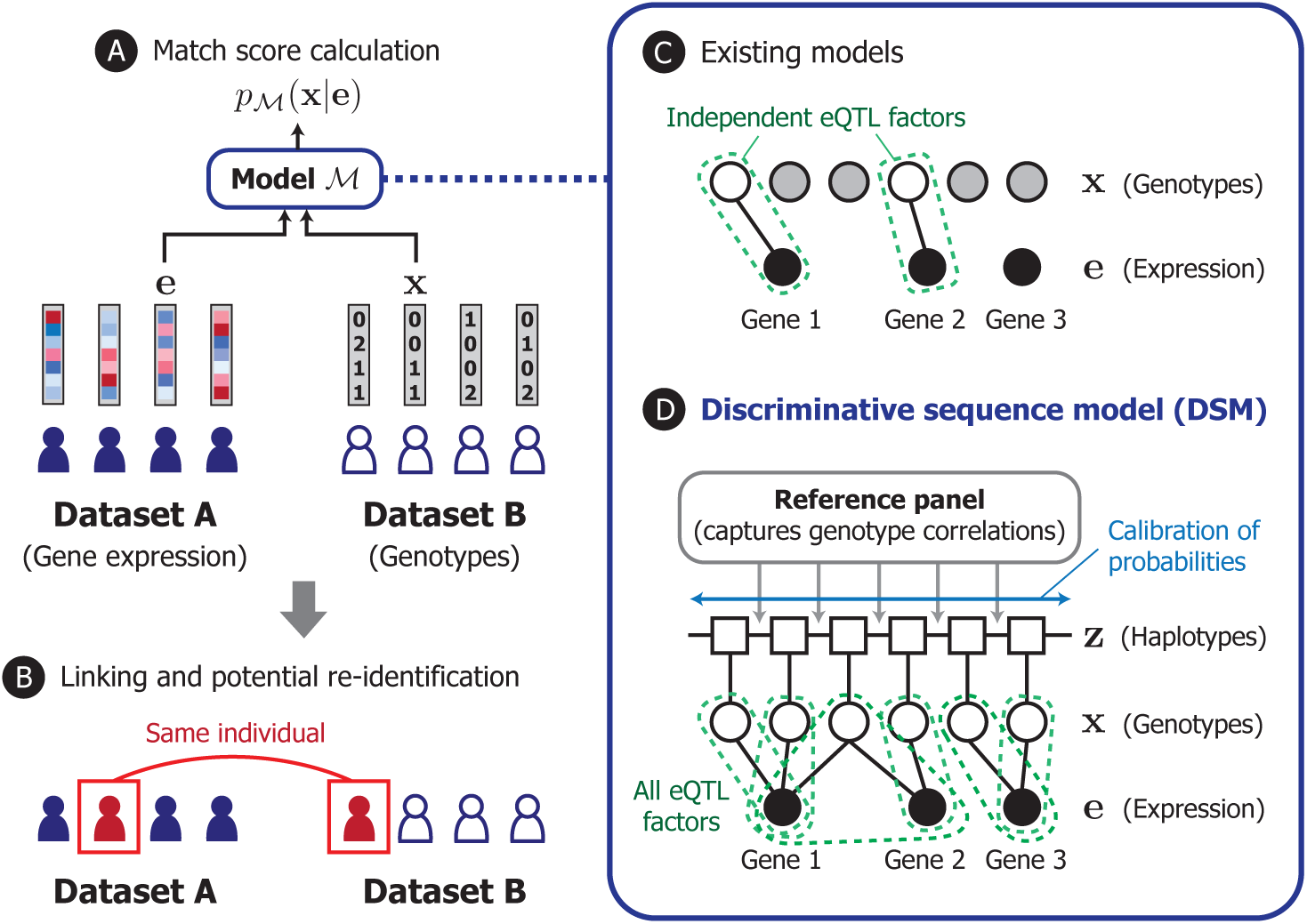
Overview of DSM. In our model, we consider the design of a match score function that quantifies how likely a given pair of gene expression and genotype profiles originated from the same individual (**A**). Such a function can be utilized by a malicious actor to link individuals across different datasets, which could lead to the re-identification of a data sample corresponding to the individual (**B**). Existing works investigating this possibility analyzed only a subset of eQTLs (i.e., unshaded nodes, genetic variants associated with gene expression levels) that are statistically independent due to model limitations (**C**). We introduce the discriminative sequence model (DSM) (**D**), which builds upon the standard Li-Stephens hidden Markov model of genetic sequences to jointly leverage the predictive signals across all known eQTLs; it incorporates necessary calibration for redundancy and correlation among eQTLs to provide a more accurate, sequence-level match score. Our modeling approach helps reveal the full extent of genotypic information in gene expression profiles to better inform privacy risk assessment.

DSM incorporates a number of novel strategies for joint modeling of genotypes and gene expression (Methods). First, instead of defining a *generative* distribution over gene expression given genotypes [Schadt et al., 2012, Harmanci and Gerstein, 2016], we take a *discriminative* approach by parameterizing the distribution over the unknown genotypes given the observed gene expression. This obviates the need for restrictive modeling assumptions over the continuous gene expression space (e.g., normality of expression levels), instead allowing us to work with discrete genotype distributions. Second, DSM learns to calibrate the predictive probabilities during training to adjust for linkage disequilibrium among nearby genetic variants as well as redundant predictive signals across different eQTLs and eGenes. This feature allows the model to leverage the full range of information captured by eQTLs without being limited to a filtered set of independent eQTLs. Third, DSM introduces eQTL probabilistic factors that are generalized to include any number of eGenes for each eQTL, rather than separately considering each eQTL-eGene pair. When considering genes with correlated expression levels, combining information across a set of genes can enhance the quality of the predictive signal. Lastly, we developed an efficient haplotype-based approximate inference scheme for DSM to enable the use of sufficiently large reference panels (which are used to capture genotype correlations) required for accurate prediction.

### Overview of Our Experiments

To evaluate the DSM, we compared the success of linking attacks based on DSM against that of two published Bayesian linking strategies, which we refer to as Gaussian Naive Bayes (GNB) [Schadt et al., 2012] and Extremity-based Linking (EBL) [Harmanci and Gerstein, 2016], described in Methods. We trained each model on pairs of genotype and gene expression profiles from the Genotype-Tissue Expression (GTEx) dataset [The GTEx Consortium, 2015], which is one of the largest available datasets for eQTL studies that could be leveraged by a potential attacker. The set of eQTLs used by the models are obtained from GTEx, representing an attacker who identifies eQTLs in the available training data. Only the reported loci of eQTLs were used; any distributional parameters were estimated during training. For DSM, we additionally used 1,000 haplotypes from the Haplotype Reference Consortium (HRC) dataset [The Haplotype Reference Consortium, 2016], excluding any overlap with GTEx, for the reference panel underlying the HMM component of DSM (Methods).

We then used the trained models to attempt a linking attack on a separate, non-overlapping dataset including both genotype and gene expression profiles, shuffled to obscure the links between the two data types. Our primary evaluation setting emulates an attack scenario where the attacker holds a gene expression profile of interest and tries to identify a genotype profile from the same individual from a pool of candidate genotype profiles in another dataset. To this end, we independently assigned the best matching genotype profile to each gene expression profile using each method and measured the overall accuracy of the attack by the number of individuals for whom we could correctly link their gene expression profiles to their genotypes. As we show below, we considered a range of attack scenarios and different choices of the candidate set to assess the robustness of our method. We evaluated the methods on Chromosomes 19 to 22.

Note that, after observing EBL’s poor performance in our experimental setting (due to relatively few SNPs per chromosome satisfying EBL’s extremity thresholds; see Methods), we instead analyzed our extension of EBL where EBL collapses to GNB for genes with non-extreme expression values. We observed that this hybrid model outperforms the original EBL model which relies only on predicted genotypes based on extreme gene expression values (see Supplemental Fig. S1 in Supplemental Methods). Hyperparameters for EBL and GNB such as the extremity threshold and eQTL-eGene correlation threshold are optimized via a grid search, and the optimal choice of parameters was used to consider the best-case performance for each method.

### DSM Links More Individuals Than Previous Approaches

We first considered the setting where the training and test datasets are both sampled from the same dataset cohort. Such a scenario may occur when an attacker obtains access to training data consisting of individuals from the same population as the target individuals. For this analysis, we used a subset of gene expression and genotype profile pairs of 450 individuals in the GTEx muscle tissue dataset [The GTEx Consortium, 2015] for training, and tested the linking accuracy on a non-overlapping, held-out set of 138 individuals also from GTEx.

DSM was able to link all 138 test individuals using all four chromosomes and 135 using Chromosome 19 alone (Table 1). Although GNB and EBL also correctly link most individuals (136 and 134 out of 138, respectively) using all four chromosomes, when using a single chromosome DSM’s linking accuracy is greater for all but one chromosome tested (22), suggesting that DSM is able to make better use of limited predictive signals. The performance gap is especially pronounced for Chromosomes 20 and 21: for Chromosome 20, GNB, EBL, and DSM correctly linked 41, 43, and 87 individuals, respectively.

**Table 1:** DSM links more individuals in GTEx cross-validation datasets. Training set and test set are sampled without overlap from the GTEx dataset. DSM links every individual in test set (138) when combining all four chromosomes, and a larger fraction of individuals on all but one chromosome individually.

| Method | Candidate set | Chr19 | Chr20 | Chr21 | Chr22 | All 4 chromosomes |
| --- | --- | --- | --- | --- | --- | --- |
| GNB | GTEx Holdout | 125 (90.6%) | 41 (29.7%) | 33 (23.9%) | 73 (52.9%) | 136 (98.6%) |
| EBL | GTEx Holdout | 126 (91.3%) | 43 (31.1%) | 34 (24.6%) | **76 (55.0%)** | 134 (97.1%) |
| **DSM** | GTEx Holdout | **135/138 (97.8%)** | **87 (63.0%)** | **65 (47.1%)** | 60 (43.5%) | **138 (100%)** |

### DSM Enhances Cross-Dataset Linking Accuracy

We next assessed our method in the setting where there is a mismatch between the populations of the training dataset and the target dataset. Here, we used all 588 pairs of gene expression and genotype profiles available in GTEx for training. For the test dataset, we used gene expression and genotype profiles of 292 individuals in the Finland-US Investigation of NIDDM Genetics (FUSION) dataset [Ghosh et al., 2000]. The gene expression data in FUSION is also obtained from muscle tissue. Notably, there is some ancestry mismatch between GTEx and FUSION, since individuals in the GTEx dataset were recruited in the United States and thus expected to have limited representation of Scandinavian ancestry, in contrast to FUSION, which includes only Finnish individuals. We used the same eQTL set and the HRC reference panel for DSM as the previous analysis, except the latter now excludes any overlap with both GTEx and FUSION.

Using DSM, we were able to link 289 of 292 individuals from Chromosome 19 alone (Table 2). Compared to GNB and EBL, our improvement is most clear on chromosomes with fewer eQTLs, such as Chromosome 21 (fewest eQTLs of four chromosomes, including around 11K), where the DSM correctly links 110 individuals while GNB and EBL link 34 and 36 individuals, respectively. All three models are able to link more than 99% of individuals when combining all four chromosomes, but the DSM provides greater linking accuracy on all four chromosomes individually.

**Table 2:** DSM enhances linking accuracy in a test population different from the training dataset. When linking individuals in the FUSION dataset with models trained on the GTEx dataset, we are able to link more individuals than previous methods, with the exception of one chromosome. When we add an additional 292 target genomes from the GOT2D consortium to the candidate set (FUSION+GOT2D), our accuracy improvement becomes more pronounced.

| Method | Candidate set | Chr19 | Chr20 | Chr21 | Chr22 | All 4 chromosomes |
| --- | --- | --- | --- | --- | --- | --- |
| GNB | FUSION | 247 (84.6%) | 74 (25.3%) | 34 (11.6%) | 162 (55.5%) | 290 (99.3%) |
| EBL | FUSION | 255 (87.3%) | 80 (27.4%) | 36 (12.3%) | 166 (56.8%) | 290 (99.3%) |
| **DSM** | **FUSION** | **289/292 (99.0%)** | **188 (64.4%)** | **110 (37.7%)** | **241 (82.5%)** | **292 (100%)** |
| GNB | FUSION+GOT2D | 224 (76.7%) | 56 (19.2%) | 17 (5.8%) | 99 (33.9%) | 286 (97.9%) |
| EBL | FUSION+GOT2D | 234 (80.1%) | 63 (21.6%) | 16 (5.5%) | 111 (38.0%) | 289 (99.0%) |
| **DSM** | **FUSION+GOT2D** | **280/292 (95.9%)** | **88 (30.1%)** | **52 (17.8%)** | **143 (49.0%)** | **292 (100%)** |

To evaluate the ability of each method to distinguish the true match from a larger set of candidates, we next added 292 additional genotype profiles (584 phased haplotypes) from the GOT2D consortium data [Flannick et al., 2019], of which FUSION is a subset, to the set of candidate genotype profiles for linking. In this scenario, we expect the set of individuals still correctly linked to be a subset of those linked in the previous experiment based only on FUSION, since increasing the number of candidates only makes the match score of the true link less likely to be the best score.

Our results show that DSM is still able to link all 292 individuals combining the four chromosomes(Table 2; Fig. 2). The accuracy of GNB and EBL was substantially reduced by the additional candidates, for example resulting in 17 and 16 correctly linked individuals, respectively, for Chromosome 21 compared to 52 by DSM. This drop in linking accuracy is observed even for a modest number of added samples (Fig. 2). With just 100 additional samples, GNB and EBL newly miss at least 10 individuals on three of the four chromosomes.

**Figure 2:**
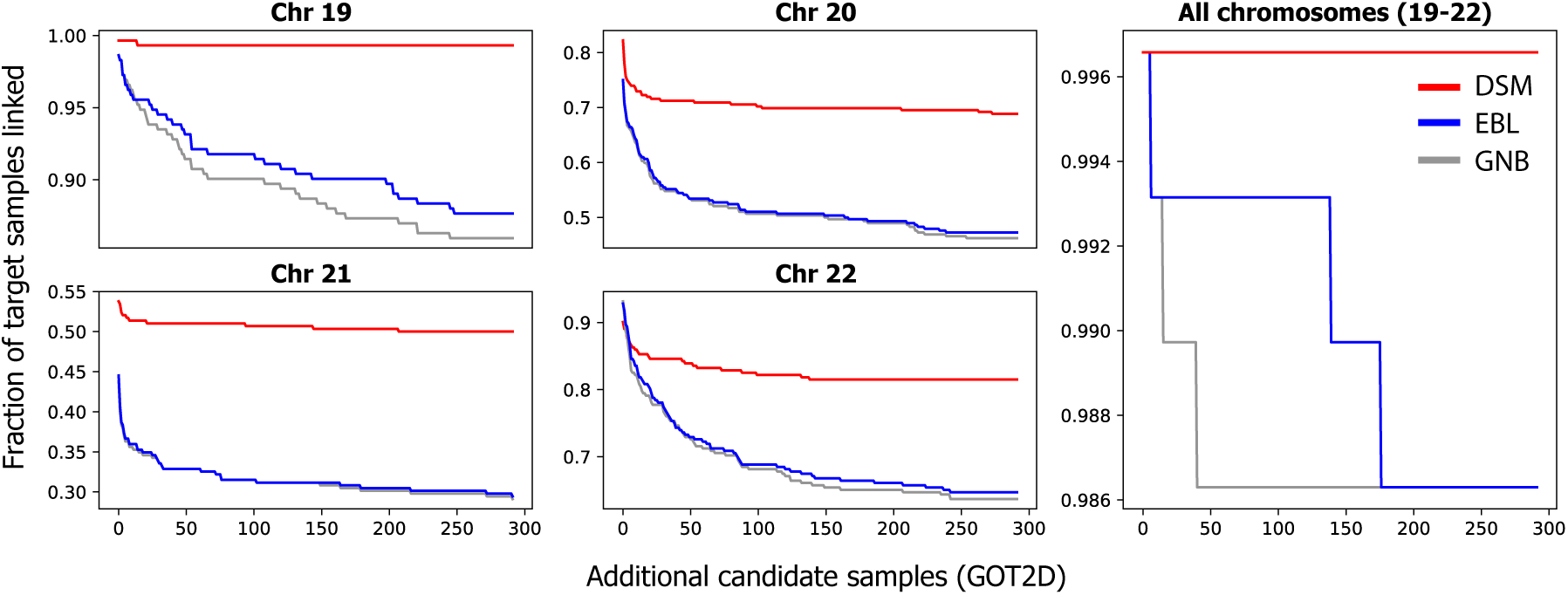
Linking based on DSM remains accurate with larger candidate genotype sets. The plots depict how the linking accuracy of each method for FUSION individuals changes as more non-matching candidate target genotypes from the GOT2D consortium are added. Results are shown for each of Chromosomes 19-22 as well as all four chromosomes combined to demonstrate performance on different sets of eQTLs. We add the same set of additional candidate samples for each chromosome. Each step corresponds to one individual unlinked. DSM remains accurate for most target individuals, even as more candidate genotype profiles are added.

### DSM Extracts Stronger Identifying Signals from Gene Expression

To gain a deeper insight into DSM’s robust performance on larger sets of candidate genotype profiles, we aimed to assess the *gap* in the match score between the true match and the remaining mismatching pairs. Intuitively, if the gap is large, then we should expect the model to remain accurate for longer as additional samples are added to the candidate set, as it would be less likely that random samples of other individuals’ genotype profiles will result in match scores as extreme as the true match.

To assess this gap, we modeled the null distribution of match scores for each of the three models (see Supplemental Note S1 in Supplemental Methods), which allowed us to compute the *p*-value representing the strength of identifying signal of each matching **x**^(*i*)^ and **e**^(*i*)^. While all three models were able to separate the matching pair from the mismatching pair for more than 99% of individuals in FUSION (Table 2) with low *p*-values, DSM was able to generate a lower *p*-value for 229 (78.4%) and 239 (81.8%) individuals compared to GNB and EBL, respectively, suggesting greater separation of the true match (Fig. 3A,B,C).

**Figure 3:**
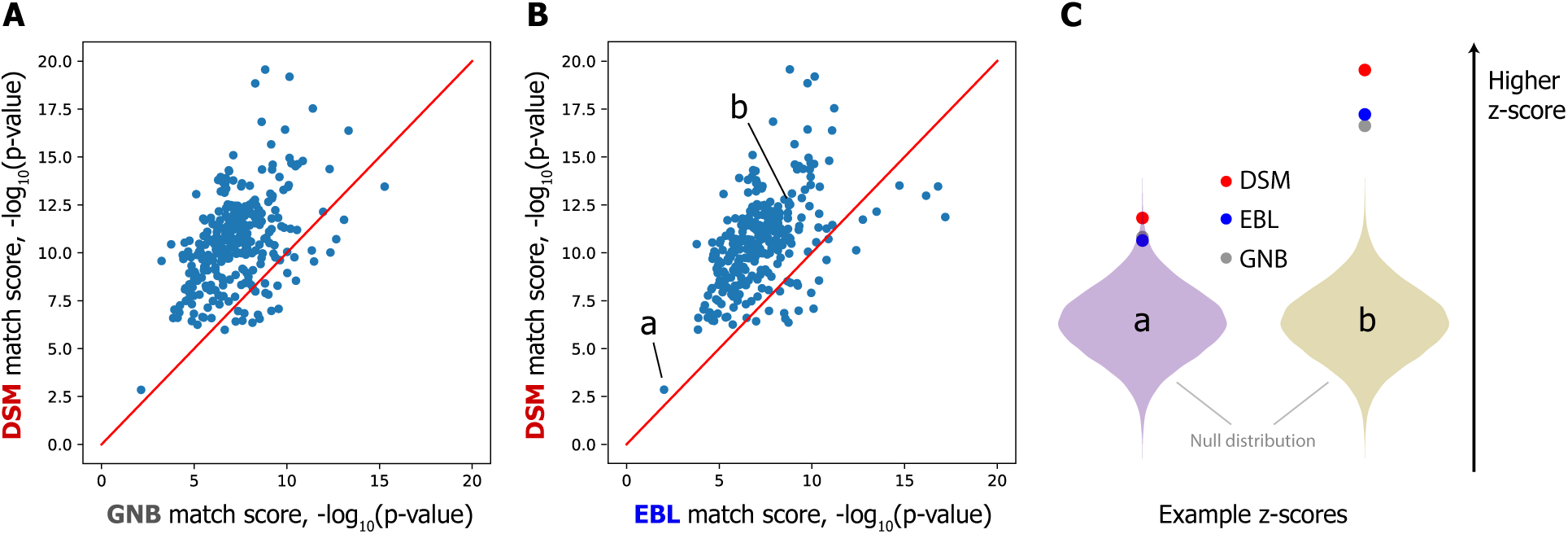
DSM distinguishes matching pairs of profiles from mismatching pairs with higher confidence. (**A** and **B**) DSM separates most individuals in the test dataset from the background distribution of match scores more distinctly than both GNB (A) and EBL (B), as assessed by the *p*-value of the true match score compared to null distribution of mismatching pairs’ scores (FUSION individuals only). In **C**, we visualize the strength of separation for the two individuals marked (a and b) in relation to the null distribution normalized as a Gaussian distribution. The DSM indeed separates these individuals more than GNB and EBL. Note that (a) is the one individual that all three methods fail to link, and this individual is substantially less separated from the null distribution than the other individuals.

This result explains our stronger performance when adding extraneous GOT2D genomes; one can imagine adding new GOT2D genomes as sampling from the null match score distribution, so if the *p*-value of the correct match is lower, more samples must be generated to find a mismatching pair with a better score than the true match. We observed lower *p*-values for DSM also based on a larger candidate set including GOT2D individuals (see Supplemental Fig. S2 in Supplemental Methods). These results indicate that DSM more sharply separates the matching pair from mismatching pairs.

### Finding the Needle in a Haystack: DSM Enables Linking against Massive Candidate Genotype Sets

We set out to test whether a linking attack based on the DSM is feasible even on a large-scale candidate genotype set including tens of thousands of individuals. This represents a plausible attack scenario where the attacker takes a target gene expression profile and searches against one of a growing number of large biobank-scale genomic datasets in which the individual may be known to be included. In such a scenario, a large number of non-matching genotype profiles must be distinguished from the true match in order to obtain a correct link.

To this end, we selected a total of 21,996 genotype profiles (43,992 phased haplotypes) from HRC that did not overlap with FUSION, GTEx, or the DSM reference panel and additionally included this cohort as part of the candidate set together with the FUSION individuals. We observed that DSM displays remarkable robustness to such large datasets, linking 260 of 292 FUSION individuals (89.0%), compared to just 129 (44.1%) and 147 (50.3%) by GNB and EBL respectively for Chromosome 20 (Table 3; Fig. 4B). When combining all four chromosomes, DSM linked 273 (93.5%), notably more than 267 (91.4%) and 260 (89.0%) for EBL and GNB, respectively (Table 3; Fig. 4A). Similar gaps were observed for all chromosomes individually (Supplemental Fig. S3 in Supplemental Methods).

**Figure 4:**
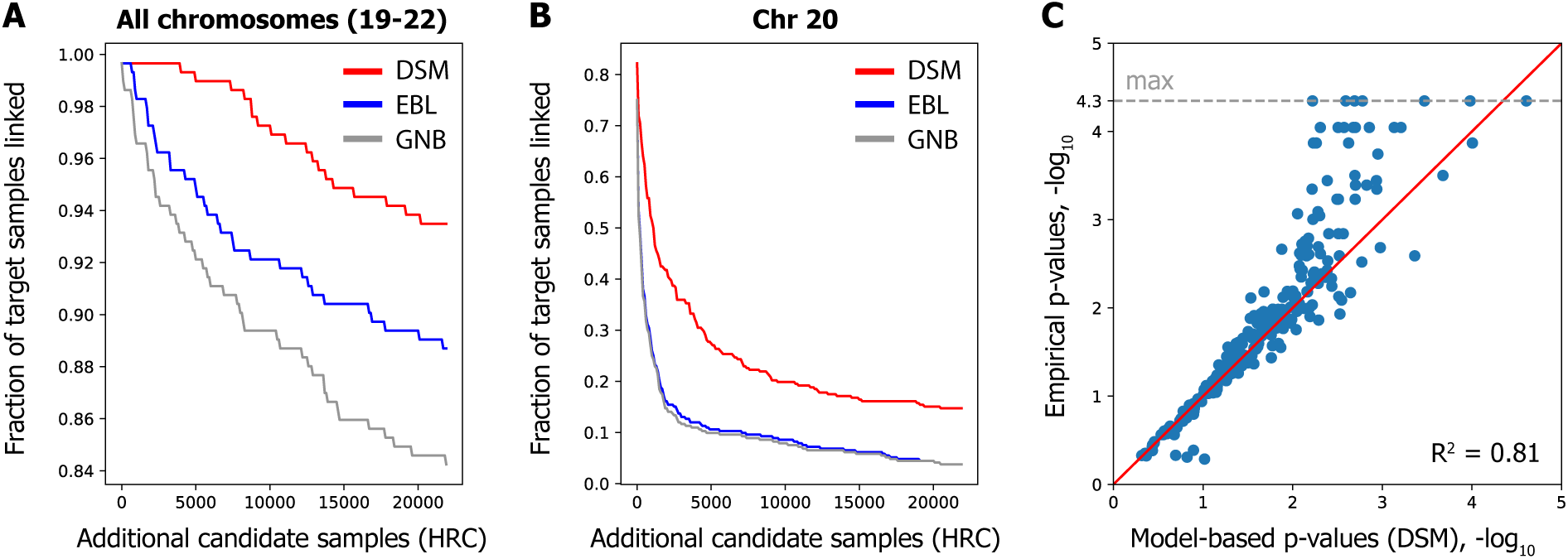
DSM enables linking on massive candidate genotype sets and provides a measure of identifying signals from gene expression data. We included an additional 22K HRC individuals in the candidate genotype set to evaluate linking accuracy on massive datasets. (A) Combining all four chromosomes, DSM links 94% of individuals. Each curve represents an average across ten random permutations of the HRC individuals. (B) DSM’s substantial accuracy improvement is observed for each chromosome, as depicted here for Chromosome 20. Plots for other chromosomes are provided in Supplemental Fig. S3 in Supplemental Methods. (C) A strong correlation is observed between the empirical *p*-values of true matches and our model-based estimates on a random subset of 500 individuals, suggesting that our *p*-values provide a calibrated measure for assessing our method’s accuracy on larger candidate sets. The minimum empirical *p*-value (1*/*22192), or equivalently maximum log *p*-value, is highlighted with the dashed gray line.

**Table 3:**
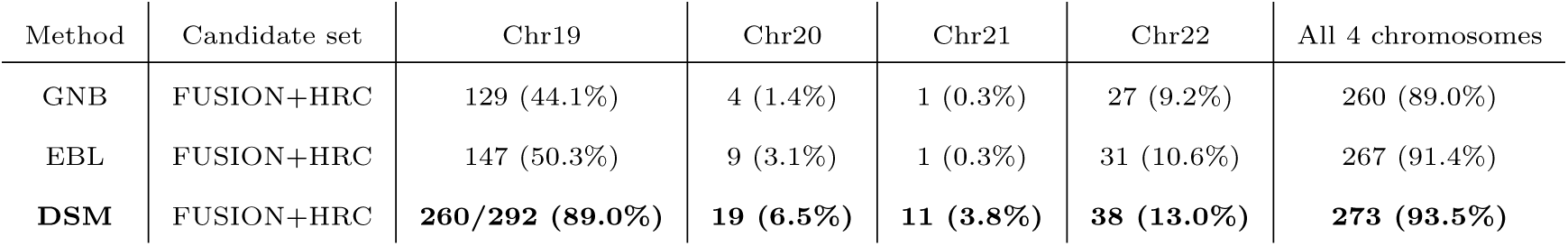
DSM robustly links individuals in massive candidate genotype sets. When the expanded candidate genotype set including the HRC cohort contains approximately two orders of magnitude more individuals (22K) than the original FUSION dataset (292), DSM consistently links more individuals than EBL and GNB across all four chromosomes.

| Method | Candidate set | Chr19 | Chr20 | Chr21 | Chr22 | All 4 chromosomes |
| --- | --- | --- | --- | --- | --- | --- |
| GNB | FUSION+HRC | 129 (44.1%) | 4 (1.4%) | 1 (0.3%) | 27 (9.2%) | 260 (89.0%) |
| EBL | FUSION+HRC | 147 (50.3%) | 9 (3.1%) | 1 (0.3%) | 31 (10.6%) | 267 (91.4%) |
| **DSM** | FUSION+HRC | **260/292 (89.0%)** | **19 (6.5%)** | **11 (3.8%)** | **38 (13.0%)** | **273 (93.5%)** |

Furthermore, testing the linking attack on a much larger candidate set allowed us to validate our model-based *p*-values, calculated on a subsampled candidate set (including 500 individuals) using our model of null distribution (Supplemental Note S1 in Supplemental Methods), by comparing them to the empirical *p*-values calculated on the full candidate set (Fig. 4C). Increasing noise is expected at the tail of the distribution given the large sample size requirements for empirically estimating small *p*-values. Nevertheless, the high concordance between the model-based and empirical *p*-values (Pearson *R*^2^ of 0.89, log-transformed) provide further evidence that the enhanced identifying signals obtained by DSM (as measured by the *p*-values) truly reflect more robust linking performance in general.

### Additional Linking Attack Scenarios

We thus far have evaluated the accuracy of linking attacks when the genotype profile of the same individual as the query gene expression profile is included in the target dataset. In an attack scenario where the membership of the matching individual in the target dataset is unknown to the attacker, the attacker must make an additional decision about whether to draw a link between an expression profile and the top matching genotype profile. For instance, this can be achieved by imposing a threshold on the match scores such that only the high-confidence links above the threshold are called. To assess the performance of DSM in this setting, we performed a holdout experiment based on the expanded FUSION dataset (with 22K candidate genotype profiles), where the matching genotypes of half of the query expression profiles (146 out of 292) were excluded from the target set before the linking procedure. For each method, we evaluated the linking results across a range of match score thresholds with respect to the standard performance metrics for binary classification, such as false positive rate (FPR), precision, and recall, which we adapted for the linking problem (see Supplemental Note S2 in Supplemental Methods).

Our results show that linking based on DSM provides a significantly better tradeoff between identifying true matches and minimizing false positives compared to EBL and GNB (Fig. 5). For example, we observed that the false positive rate (FPR; the proportion of query expression profiles without any matching genotypes that were incorrectly linked) of our linking approach is small (*<*1%) when identifying half of the true matches with top match scores, indicated by a true positive rate (TPR) of 50% (Fig. 5A). Naturally, identifying more true matches leads to a higher FPR (e.g., a FPR of 8% for a TPR of 80%). In contrast, EBL and GNB led to substantially higher FPRs for the same TPR; e.g., for a TPR of 80%, they obtained a FPR of 35% and 34%, respectively. In agreement with these observations, DSM obtained a greater AUROC metric (0.887) than both EBL (0.809) and GNB (0.799). We observed a similar improvement of DSM in terms of the tradeoff between the precision and recall metrics across different match score thresholds (Fig. 5B).

**Figure 5:**
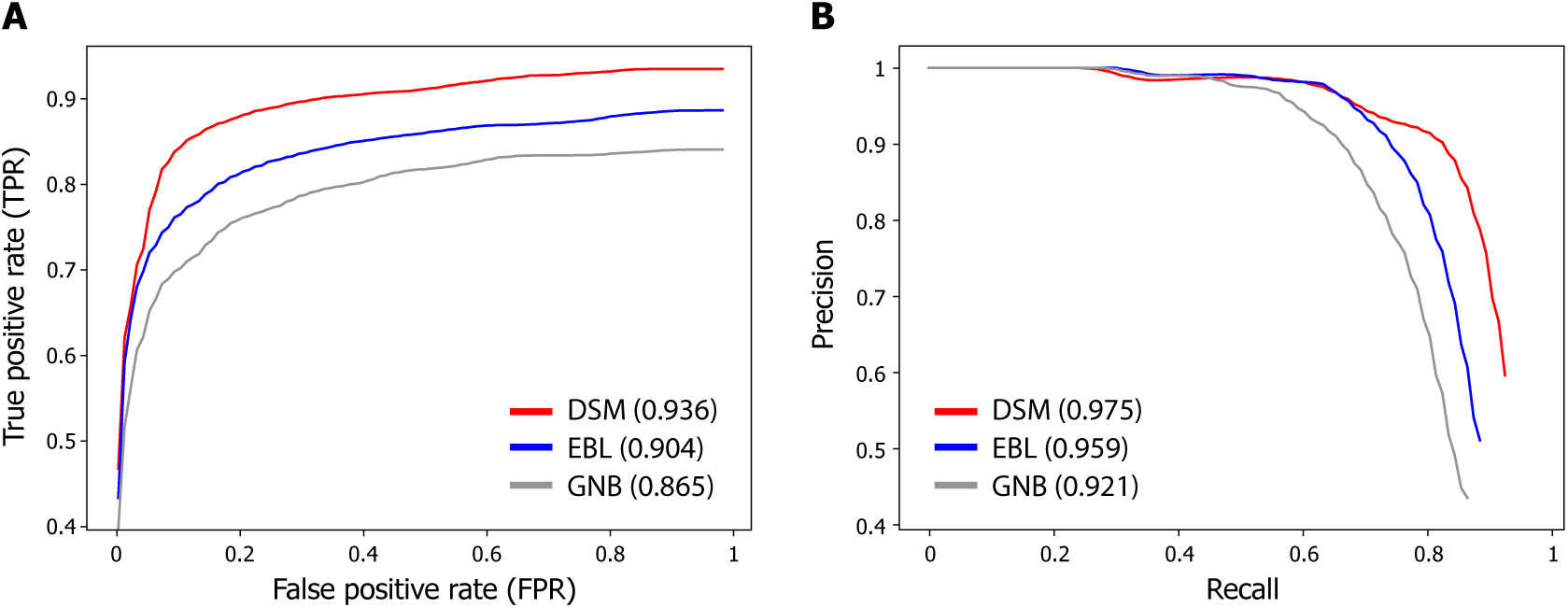
DSM leads to fewer false positives when linking individuals with unknown membership in the target genotype set. When it is unknown whether the individual associated with a query expression profile is included in the target dataset, a match score threshold can be used to draw links only for scores greater than the threshold. The choice of this threshold determines the tradeoff between detecting true matches and avoiding spurious links. Receiver-operating-characteristic (ROC) curves (**A**) and precision-recall curves (**B**) illustrating this tradeoff are shown for DSM, EBL, and GNB, evaluated on the expanded HRC dataset with 22K individuals. The average AUROC and AUPRC metrics are reported in parentheses for each method. Full descriptions of the relevant evaluation metrics (e.g., true positive rate and false positive rate) are provided in Supplemental Note S2 in Supplemental Methods. The curves are averaged over 100 trials of the holdout experiment where the genotype profiles of a random half of FUSION individuals were excluded from the target set before each trial of the linking procedure to assess false detection error. DSM outperforms previous methods in finding more true matches while minimizing the number of incorrect links for individuals without a match.

We next investigated another relevant attack scenario where the attacker holds a genotype profile and wishes to match to a gene expression profile in the target dataset. We refer to this as *reverse* linking to distinguish it from our main attack scenario. In this setting, the attacker could directly leverage an existing method for predicting gene expression from genotypes to compare with the candidate expression profiles. To this end, we compared our method—using the same scores from DSM as before, but performing linking in reverse—against linking based on MetaXcan [Barbeira et al., 2018], a state-of-the-art method for gene expression prediction. We considered two different match scores for MetaXcan: Pearson and Spearman’s correlation coefficients between the predicted and observed expression profiles. Training the models on the GTEx muscle-skeletal dataset and evaluating reverse linking on the FUSION dataset, we observed perfect linking accuracy for DSM when combining all four chromosomes (291/292), substantially more accurate than MetaXcan-based linking (Pearson: 171/292, Spearman’s: 97/292) (see Supplemental Table S1 in Supplemental Methods). For individual chromosomes, reverse linking was generally less accurate than our original results in the forward direction based on DSM (Table 2), but DSM still obtained greater linking accuracy than MetaXcan. We hypothesize that the reduced accuracy of reverse linking is due to the greater impact of noise (both biological and experimental) in gene expression profiles on reverse linking, since a large number of expression profiles are jointly considered to determine the match for each query genotype profile.

## Discussion

We have shown that existing models for transcriptomic linking attacks overlook a substantial amount of predictive signals captured by eQTLs. Our *discriminative sequence model* (DSM) poses the linking attack as a sequence-level probabilistic modeling problem, one in which the attacker jointly predicts the entire genotype sequence instead of independently predicting each eQTL. Our results demonstrate that DSM reveals greater linking attack risk than previous methods over a range of evaluation settings, including cross-dataset prediction and linking attacks involving a massive candidate set. We also illustrated how the *p*-values calculated based on the match score distributions can be a proxy for the strength of identifying signals captured by the model, representing a promising new approach for quantifying privacy risks.

Our framework unifies QTL privacy analyses, since it allows joint prediction of genotypes from any source. It would be interesting to explore incorporating other sources of phenotypic data that could leak genotype information into the DSM, including protein abundance (pQTLs) [Bowler et al., 2022], methylation (mQTLs) [Gaunt et al., 2016, Backes et al., 2017], and allele-specific expression (aseQTLs) [Gürsoy et al., 2021]. Another promising direction would be extending the DSM formulation to the co-expression of genes conditional upon eQTLs, which has been used by other models [Gamazon et al., 2015, Gusev et al., 2016].

There are a few limitations of our work. Training the DSM requires substantial memory (75GB) and time (*∼*6 hours) per window. These requirements depend on the size of the eQTL window and reference panel size (see Supplemental Fig. S4 in Supplemental Methods). In contrast, both EBL and GNB can be trained with minimal time and memory requirements. However, once trained, DSM performs matching with far less memory (12GB) and time (around 5 minutes) than required for training, resulting in less than a day of runtime for the large-scale matching we demonstrated with 22K individuals. Also, for assessing the risk of linking attacks, it may be desirable to consider an adversary with large computational resources.

While our work reveals that gene expression data contains more identifying information than previously known, we acknowledge that further investigation is needed to ascertain the real-world implications of our findings in the context of existing transcriptomic databases. For instance, when the target genotype set does not include the individual associated with the gene expression profile, the risk of a successful linkage is reduced as only the matches with especially high match scores can be called with sufficiently low false detection error, as we illustrated in our experiments (see Additional Linking Attack Scenarios). Furthermore, there are other key factors that modulate the success of potential linking attacks, including ancestry, assay platform for gene expression measurements, and the tissue of origin of the samples. When the target set is different than the training set in these aspects, eQTL-based linking may have reduced accuracy but would still remain feasible [Harmanci and Gerstein, 2016]. Thus an appropriate level of protection for gene expression data would ultimately depend on the specific setting of the dataset as well as the availability of other public data resources that could be jointly leveraged for an attack.

Beyond standard access control mechanisms, other tools for consideration to enhance data protection include data sanitization [Gürsoy et al., 2022b, He et al., 2018, Zhang and Bonomi, 2022, Yilmaz et al., 2020, Ye et al., 2022, Harmanci and Gerstein, 2018], differential privacy [Tramèr et al., 2015, Almadhoun et al., 2020, Chen et al., 2020], and secure analysis and storage platforms [Gürsoy et al., 2022a, Dokmai et al., 2021, Froelicher et al., 2021, Cho et al., 2022, Lambert et al., 2018]. Investigating the use of these techniques to provide meaningful mitigation of the risks illustrated in our work, while maintaining the scientific utility of sharing genomic and transcriptomic data, is an important next step. All these efforts lay the foundation for secure sharing of omics data.

## Methods

### Problem Description and Threat Model

We consider the setting where a malicious actor (attacker) wishes to link an individual’s gene expression profile to his or her genotype profile in a different dataset [Schadt et al., 2012, Harmanci and Gerstein, 2016]. Both gene expression and genotype data could be either acquired from a public repository or accessed through an authorized channel. In common data sharing scenarios, both types of data are provided without explicit personal identifiers; however, they may include clinical and demographic attributes deemed necessary for the respective studies. These additional attributes, when combined between the two data sources, may lead to the re-identification of an individual even when their identity is not known to the attacker [Sweeney, 2000]. Studies have also shown that the genetic sequence by itself can be a sufficient identifier when it is searched against public genealogy databases [Gymrek et al., 2013, Erlich et al., 2018]. These findings illustrate a plausible threat of re-identification when a gene expression profile is linked to the corresponding genetic sequence.

The goal of the attacker is to utilize a match score function that, given a query gene expression profile, scores all target genotype profiles as possible links based on some criterion. We consider this function to be the attacker’s most important tool, which must be learned properly in order to successfully carry out a linking attack. We thus assume that a motivated attacker will obtain access to a few additional datasets for learning the match score function, including: (i) a publicly available list of known eQTL associations (e.g., in the GTEx Portal [The GTEx Consortium, 2015]), (ii) a training dataset of genotype and gene expression profile pairs, and (iii) a set of genetic sequences (without associated gene expression data) for modeling genotype distributions. We note that the latter two data sources are commonly available in existing data repositories (e.g., NIH dbGaP [Tryka et al., 2014]) and are generally growing in size and availability. While data use agreements for public repositories typically disallow attempts for re-identification, it is worth noting the attacker could use these additional sources only to train a match score function and not to re-identify individuals in those datasets. Moreover, without clear mechanisms for data provenance and for detecting a breach of agreements, it is plausible that an attacker could use these datasets without repercussions.

For the purpose of comparing the effectiveness of different match scores, we primarily consider the setting in which the attacker knows that a given query gene expression profile matches one of the genotype profiles in the target dataset. In practice, the attacker may not know the membership of the individual in the target dataset and thus might need to make an additional decision about whether the best possible match found truly represents the same individual (e.g., by imposing a minimum threshold on the match score). In *Additional Linking Attack Scenarios*, we also address the performance of our method in this setting as well as for the closely related problem of matching a genotype profile across a set of candidate gene expression profiles (i.e., linking in the opposite direction from that of our main setting).

### Notation and Definitions

We first provide a formal definition of the transcriptomic linking problem. The attacker has access to two datasets *D_X_* and *D_E_*, which correspond to a dataset of genotypes and gene expression profiles, respectively. We let each **x**^(*j*)^ *∈ D_X_* be a phased genotype profile over biallelic variants (a pair of haplotypes), that is, **x**^(*i*)^ *∈ {*(0, 0), (0, 1), (1, 0), (1, 1)*}^V^*, where *V* is the number of variants considered. Any unphased genotype dataset can be preprocessed using standard phasing algorithms to obtain a phased dataset (e.g., [Loh et al., 2016, Browning et al., 2021, Delaneau et al., 2019]). Each **e**^(*j*)^ *∈ D_E_* is a vector of gene expression level measurements obtained via microarray or RNA sequencing experiments, i.e., **e**^(*j*)^ *∈* R*^G^*, where *G* is the number of genes (e.g., around 20,000).

For each individual gene expression profile **e**^(*j*)^ *∈ D_E_*, the attacker scores all candidate genotype profiles **x**^(*i*)^ *∈ D_X_* according to some match score function *M* (**x**^(*i*)^, **e**^(*j*)^). This function must return a high match score if **x**^(*i*)^ and **e**^(*j*)^ are collected from the same individual, and a low score otherwise. It is natural to use a probabilistic interpretation for *M* such that *M* (**x**^(*i*)^, **e**^(*j*)^) = *p*(**x**^(^*^i^*^)^*|***e**^(^*^j^*^)^) (or *p*(**e**^(^*^j^*^)^*|***x**^(^*^i^*^)^)), since this represents the probability of observing the target genotype given the query gene expression vector (or vice versa). As previously described, we assume that the attacker trains the match score function *M* based on an independent training dataset *D^′^* = *{*(**x***^′^*^(*i*)^, **e***^′^*^(*i*)^)*}_i=1_^N^*, including matching pairs of genotype and gene expression profiles. With this formulation, obtaining a suitable *M* thus becomes a statistical learning problem, where the attacker optimizes the parameters of *M* to maximize the likelihood *p*(**x***^′^*^(^*^i^*^)^*|***e***^′^*^(^*^i^*^)^) over the training dataset, then relies on the generalization of *M* to perform linking between the target datasets *D_X_* and *D_E_*.

### Review of Previous Linking Approaches

Previous works have shown that using a learned model to define the match score function *M* can provide the ability to predict genotypes based on gene expression [Schadt et al., 2012, Harmanci and Gerstein, 2016]. These works leverage a set of genetic variants, called eQTLs, whose genotype values are correlated with the expression levels of a gene and thus can be predicted given a gene expression profile.

#### Gaussian naïve Bayes (GNB) approach [Schadt et al., 2012]

From here onwards, **x** (of length *V*) refers only to the eQTL subset of the genotype profile. Also, let *Q*(*s*) *⊂* [*G*] denote the set of genes associated with eQTL *s*, and **e***_Q_*_(*s*)_ denote the corresponding subset of **e**. Schadt et al. [Schadt et al., 2012] begin by observing

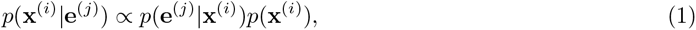

by Bayes’ theorem. To simplify the prediction, they further make an *independence* assumption across eQTLs to express *p*(**x**^(^*^i^*^)^*|***e**^(^*^j^*^)^) as a product of probabilities for individual eQTLs:

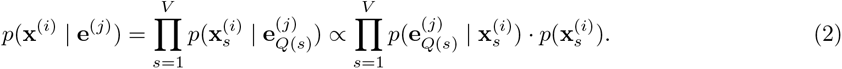

The final expression is used as the match score with additional heuristics for normalization. Both probability terms in the score are estimated on the training data: The conditional distribution over gene expression *p*(**e***_Q_*_(_*_s_*_)_*|***x***_s_*) (with a singleton *Q*(*s*) in the setting of Schadt et al.) is parameterized as a Gaussian distribution *N* (*µ_s_, σ_s_*) with mean *µ_s_* and standard deviation *σ_s_*, whereas the prior distribution *p*(**x***_s_*^(*i*)^) is determined by the genotype frequencies. We refer to this approach as Gaussian näıve Bayes (GNB), given the Gaussian conditional distribution and the factorization based on independence assumption, which is analogous to näıve Bayes classifiers.

#### Extremity-based linking (EBL) approach [Harmanci and Gerstein, 2016]

Given the noisy nature of gene expression measurements, the strongest predictive signals often lie in the extreme values of gene expression levels, where a Gaussian model may not lead to reliable probability estimates. Harmanci and Gerstein [Harmanci and Gerstein, 2016] leverage this insight to develop an alternative linking strategy that directly utilizes extreme gene expression values for prediction. Specifically, they set 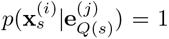 (considering only singleton *Q*(*s*)), for **x_s_**^(*i*)^ = (0, 0) or (1, 1) depending on the direction of association, when (i) 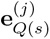 is sufficiently “extreme” according to its empirical distribution, and (ii) **x***_j_* and **e***_Q_*_(*s*)_ are strongly correlated, for suitable thresholds for extremity and correlation. The match score is then computed by comparing the predicted genotypes and the given candidate genotype profile.

Importantly, both models treat each SNP independently. To ensure that this assumption is justifiable, the set of eQTLs used by previous models are pruned to a smaller set of mutually independent eQTLs. However, this approach omits all other eQTLs which collectively provide far richer information about the genotypes than a pruned set would provide, as we show in our work. Simply including all eQTLs in the prediction, disregarding the correlations and redundant predictive signals, lead to poor predictive performance due to miscalibrated probabilities (see Supplemental Fig. S5 in Supplemental Methods), which necessitates our new modeling approaches.

Other works have explored re-identification attacks based on sparse and noisy genotypes (e.g., those obtained from a coffee cup), where a hidden Markov model (HMM) is used to infer matches while considering the haplotype structure [Emani et al., 2021]. These works leverage the key insight that higher-order correlations (LD) exist between SNPs, and that recombination modeling enables more accurate privacy risk estimation in these scenarios [Samani et al., 2015, Deznabi et al., 2017]. While our work is motivated by a similar insight, unlike the existing works, we introduce an end-to-end learning framework for sequence-level prediction of genotypes based on weak statistical signals from non-genotypic data and demonstrate the effectiveness of this approach on gene expression data. Furthermore, acquiring calibrated probabilities for target genotypes is a challenging yet important task for risk quantification and is uniquely addressed by our work.

### Our Novel Approaches for Predicting Genotypes from Gene Expression

Here, we summarize the modeling techniques we newly leverage to improve upon the existing models of genotype prediction based on gene expression.

#### Discriminative probabilistic modeling

Recall that our goal is to learn a match score function *M* (**x**, **e**) in terms of *p*(**x***|***e**) to obtain a ranking over candidate genotype profiles **x** given a particular expression profile vector **e**. As we described, existing methods use the relation *p*(**x***|***e**) *∝ p*(**e***|***x**)*p*(**x**) to instead model the conditional distribution over expression given genotypes. This is considered a **generative** approach to probabilistic modeling, where the prediction is made by hypothesizing different values of the unobserved variable (**x**) and choosing one that most likely have generated the observed data (**e**). However, since **e** is high-dimensional and has a continuous domain, any parameterization of *p*(**e***|***x**) must make strong assumptions about its distribution (e.g., normality), which most likely introduces inaccuracies. Also, when multiple genes are associated with an eQTL, modeling the joint distribution *p*(**e***_Q_*_(_*_s_*_)_*|***x***_s_*) is expected to be even more challenging; this in part explains why previous works only considered single-gene settings. Instead, we adopt a **discriminative** approach, whereby the target distribution *p*(**x***|***e**) is directly parameterized and learned. This obivates the need to model the distribution over **e**, significantly simplifying the problem. In addition, **x** is a vector of discrete values, which requires fewer parametric assumptions.

#### Capturing genotype correlations

Previous models assumed that *p*(**x***|***e**) could be factorized into independent probabilities for each eQTL, i.e., ∏*_s_ p*(**x***_s_|***e***_Q_*_(_*_s_*_)_), which allows each term to be estimated and calculated independently. However, this requires pruning of correlated eQTLs, resulting in a smaller set of eQTLs to use for prediction and obscuring the true extent of genotypic information leakage. To address this challenge, we incorporate the Li and Stephens’ model of haplotype sequences [Li and Stephens, 2003], based on a hidden Markov model (HMM), as the backbone of our probabilistic model for *p*(**x***|***e**). Note that the HMM defines a probability distribution over a haplotype sequence, viewing it as a mosaic copy of a collection of haplotype sequences in a reference panel. At each genomic position, there is a probability of crossing over to a different reference haplotype to copy from, which represents the recombination process. This model component thus captures the correlation (i.e., linkage disequilibrium) structure among nearby eQTLs, allowing us to make predictions that correctly account for this structure while considering all eQTLs.

#### Going beyond single-gene predictions

Another shortcoming of previous models is that they could only consider single eQTL-eGene pairs at a time, i.e., *p*(**e***_Q_*_(_*_s_*_)_*|***x***_s_*) with a singleton *Q*(*s*). In reality, multiple genes can be associated with a particular eQTL and the full set of associated genes may provide richer and more accurate information about the genetic variant than only considering the most significant gene. For example, one could reduce the experimental noise in gene expression data by averaging information across correlated genes. Our model allows any number of genes to be used in calculating the predictive signal for a particular eQTL, captured by the following probabilistic factor in our model, instantiated for each eQTL *s*:

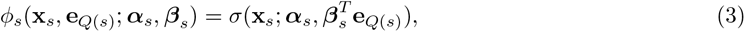

This component allows us to include an arbitrary number of genes in the partial prediction of **x***_s_* (further refined through the global model) without overly increasing model complexity.

### Discriminative Sequence Model (DSM)

We newly introduce DSM, a probabilistic graphical model that enables joint prediction of **x** from **e** by combining the insights described above. More precisely, we model the conditional distribution *p*(**x***|***e**) as a conditional random field (CRF) [Sutton et al., 2012], including probabilistic factors capturing genotype correlations as well as genotype-expression associations given by the eQTLs. A graphical representation of DSM is provided in Supplemental Fig. S6 in Supplemental Methods.

We denote the two haplotypes of **x** as **h***^m^* (maternal) and **h***^p^* (paternal). DSM models the prior over each haplotype using the Li and Stephens’ HMM [Li and Stephens, 2003] [Rubinacci et al., 2021] with respect to a given reference panel. The HMM includes a chain of hidden states **z***^m^* and **z***^p^* (for **h***^m^* and **h***^p^*, respectively), which indicate, at each position, the index of reference panel haplotype from which the observed genotype is copied with a small probability of error; this copying distribution is captured by an emission factor, *ɛ*(*h_s_|z_s_*), which relates the probability of the observed genotype at position *s* (*h_s_*) conditioned on the indexed reference panel haplotype at that position (*z_s_*). Each adjacent pair of hidden states are related through a transition factor, *τ* (*z_s_|z_s−_*_1_), which defines the probability of crossover between the haplotypes at positions *s−* 1 and *s* based on position-specific recombination rates [Loh et al., 2016]. In addition to these two groups of factors that make up the pair of HMMs, DSM introduces a QTL factor *ϕ_s_* for each eQTL *s*, capturing the dependence between the eQTL genotype and the expression levels of one or more genes, as defined in Eq. 3.

The full joint probability distribution (conditional and unnormalized) can be written as

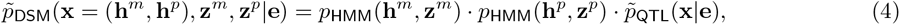

where

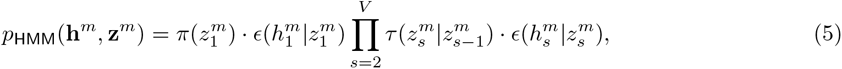

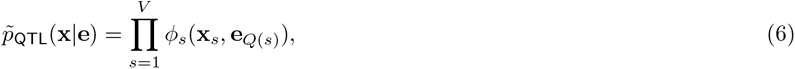

and *p*_HMM_(**h***^p^,* **z***^p^*) defined analogously to *p*_HMM_(**h***^m^,* **z***^m^*). *π* represents the prior distribution over the first hidden state, typically a flat (uniform) prior.

To apply the model, we wish to compute the marginal distribution over **x** to use for the match score:

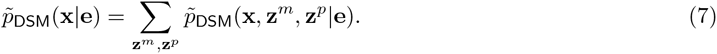

We observe that this integration can be computed efficiently using an extension of the forward-backward algorithm commonly used for inference over HMMs [Rubinacci et al., 2021, Browning and Browning, 2007, Collins, 2013]. This is achieved by considering each *ϕ_s_* as a noisy observation of the corresponding genotype variables *h_s_^m^* and *h_s_^p^*, and incorporating them into the sequential belief updates along with the emission factors in the original algorithm. We provide the details of this algorithm in Supplemental Note S3 in Supplemental Methods.

#### Haplotype approximation

Given the quadratic state space at each position when jointly considering **z***^m^* and **z***^p^*, the aforementioned inference procedure can be computationally expensive given a large reference panel. Using a large panel is necessary to accurately model the linkage disequilibrium patterns. Thus, we introduce the following modification of the QTL factors:

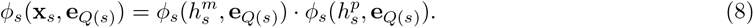

Intuitively, this approximates the predictive signal with a haplotype-specific effect that is equally applied to both haplotypes of the genotype profile. We then define

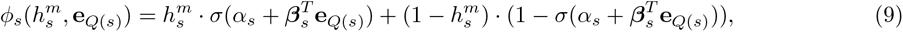

and analogously for *ϕ_s_*(*h^p^,* **e***_Q_*_(*s*)_), where *σ* denotes the sigmoid function. The learnable parameters *α_s_* and ***β****_s_* are shared between the two haplotypes. The consequence of this modification is that now we have the factorization

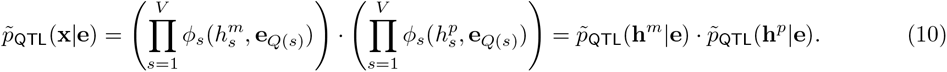

This fully decouples the two haplotypes and allows us to express

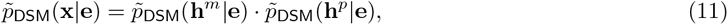

where each term is computed using the aforementioned forward-backward algorithm at the haplotype level. This reduces the overall runtime and memory requirements from quadratic to linear in the number of reference haplotypes, enabling us to leverage larger reference panels. Similarly, we obtain a memory requirement linear in the number of eQTLs considered (see Supplemental Fig. S4 in Supplemental Methods).

#### Joint learning of QTL factors

DSM provides a way to compute the probability of observing genotypes **x** given an expression profile **e**, where the adjustments informed by eQTLs are encoded by the *ϕ* factors. To ensure that the *ϕ* factors are calibrated with respect to redundant predictive signals and genotype correlations, we jointly learn all *ϕ* factors to directly maximize *p*_DSM_(**x***^′^*^(^*^i^*^)^*|***e***^′^*^(^*^i^*^)^) across all matching pairs (**x***^′^*^(*i*)^, **e***^′^*^(*i*)^) in the training dataset. This is in contrast to splitting the learning procedure to first separately training *ϕ_s_* for each *s* to predict individual eQTLs and then using these predictions to compute a sequence-wide matching score, an approach followed by all of the existing works to our knowledge [Harmanci and Gerstein, 2016, Schadt et al., 2012]. Neglecting the dependence among different eQTLs while combining their predictions can lead to miscalibrated scores with poor predictive performance (see Supplemental Fig. S5 in Supplemental Methods).

#### Match score function

For evaluating the linking performance of DSM, we use the following match score:

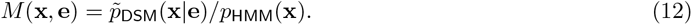

The rationale for normalizing the output of DSM by *p*_HMM_(**x**) is as follows. Since we take a particular expression profile **e** and match it across candidate **x**’s, *p*(**e***|***x**) is our desired choice for the score so as not to bias the match toward **x**’s that are more likely just based on the prior. Note that *p*(**e***|***x**) = *p*_DSM_(**x***|***e**)*p*(**e**)*/p*_HMM_(**x**), and since *p*(**e**) and the normalization factor for *p̃*_DSM_(**x***|***e**) are constant given a fixed **e**, this is equivalent to matching using the above score *M*.

### Implementation details

We implemented the forward-backward inference algorithm for DSMs in PyTorch [Collins, 2013], leveraging the automatic differentiation features and the Adam optimizer for parameter learning. We set our learning rate to 0.025 and number of epochs to 50 based on cross-validation, which was performed on a sample window of 750 eQTLs and by holding out a subset of 58 GTEx individuals (out of 588) for validation. Selected hyperparameters were used on the full GTEx dataset for all our experiments. For parallelization, we trained a separate DSM for each genomic window of 750 eQTLs with 75GB RAM and 4 CPUs for a runtime of around 8 hours. Evaluation of runtime and memory scaling with respect to window size and reference panel size is provided (see Supplemental Fig. S4 in Supplemental Methods).

For the baseline methods (GNB and EBL), we implemented the pruning of eQTLs by greedily selecting the most significant eQTL (as reported in the training dataset), then removing any other eQTLs that are correlated with it. The choice of correlation threshold was a hyperparameter we optimized, and we found that removing eQTLs with correlation greater than 0.1 to the significant eQTL consistently led to best linking accuracy on FUSION individuals. We thus adopted this threshold for comparison with our approach. Correlation was calculated as the Pearson correlation coefficient of observed genotypes between each pair of eQTLs in the training set.

## Data sets

We obtained our datasets through NIH NCBI dbGaP [Tryka et al., 2014] and the European Genome-Phenome Archive (EGA) [Lappalainen et al., 2015]. From dbGaP, we downloaded GTEx v8 muscle tissue expression (phe000037.v1) and genotype (phg001219.v1) datasets [The GTEx Consortium, 2015]; and FUSION expression (phe000033.v1) and imputed genotype (phg001194.v1) datasets [Ghosh et al., 2000]. We downloaded the HRC reference panel from the EGA (EGAS00001001710) [The Haplotype Reference Consortium, 2016]. Genotype samples in our datasets that were not already phased were phased with the Michigan Imputation Server [Das et al., 2016] using the Eagle2 software [Loh et al., 2016]. Gene expression profiles are normalized with PEER factor normalization using the default parameter setting [Stegle et al., 2012].

We included in our models all eQTL associations reported in the GTEx dataset that overlapped with the genotype data in FUSION, which consisted of: 47,322 eQTLs and 735 eGenes for Chromosome 19; 21,685 eQTLs and 254 eGenes for Chromosome 20; 11,740 eQTLs and 118 eGenes for Chromosome 21; 19,241 eQTLs and 264 eGenes for Chromosome 22; and 99,988 eQTLs and 1,011 eGenes for all four chromosomes combined.

## Software availability

Our Python implementations of DSM training and linking algorithms and example data formats and scripts are provided as Supplemental Code and also available at: https://github.com/shuvom-s/DSM.

## Acknowledgements

S.S., D.F., and H.C. conceived the project. S.S. and H.C. designed the experiments and developed the method. S.S. and D.F. implemented the software and performed the experiments. B.B. and H.C. supervised the project. All authors analyzed the data and wrote the manuscript. This work is accepted for an oral presentation at RECOMB 2023. This work is supported by the Hertz Fellowship and NSF Graduate Research Fellowship under grant number 2141064 (to S.S.); NIH R01 HG010959 (to B.B.); and NIH DP5 OD029574 and RM1 HG011558 (to H.C.).

## Supplemental Note 1: Model-based estimation of p-values

Let **M** = [*M*_1_*, · · ·, M_N_*] be a sorted vector of match scores calculated for a particular individual, and let *M^∗^* be the score for the true match. When the true match score is the largest match *M^∗^* = *M_N_*, then the individual is correctly linked. In this case, we wish to calculate a measure of how strong this identifying signal is based on the relative magnitude of *M^∗^* compared to the rest of the match scores in **M**. We achieve this by estimating a *p*-value based on a parameterized null distribution that is fit to the non-matching samples in **M**. We use a Gaussian distribution to model the null distribution, after empirically observing the approximate normality of the null distributions.

For robust estimation of the model parameters, we trim both tails of the empirical distribution (also excluding the true match) before estimating the mean and the variance of the distribution. For instance, the mean is estimated as

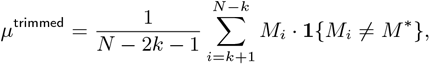

where **1***{·}* is an indicator function, and *k* determines the amount of trimming (*k/N* = 0.2 in our experiments). The standard deviation *σ*^trimmed^ is similarly estimated using this trimmed distribution, then scaled to obtain an unbiased estimator using the theoretical quantity from the corresponding truncated Gaussian distribution.

The *z*-score of our true match *M^∗^* is thus

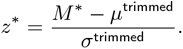

The *p*-value is calculated by first taking the tail probability of *z^∗^* in the standard normal distribution. We then re-weight the *p*-value obtained from the *z*-score by multiplying by a weight *w*, defined as

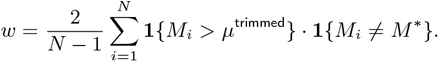

This represents two times the proportion of match scores that lie above the trimmed mean, which corrects for potential asymmetry in the null distribution below and above the mean. Empirically, we observe that our above approach leads to accurate model-based *p*-values that are consistent with permutation-based, empirical *p*-values (see Fig. 4C).

## Supplemental Note 2: Evaluation metrics for linking with match score thresholds

To investigate an attack scenario in which the attacker must make a decision about whether the proposed match is a true match, we evaluated the full range of match score thresholds for each of the three methods with respect to a number of standard metrics for binary classification. We extended these metrics to the setting of linking attacks. The key difference in our setting is that some data instances may never be linked correctly regardless of the threshold, if an incorrect match with a higher match score exists.

For each trial of our holdout experiment, we randomly split the individuals in our test dataset (FUSION) into two halves, which we term Set 1 and Set 2. We keep the genotype profiles of Set 1 individuals in the target genotype set, while excluding them for Set 2 individuals in order to assess the rate of false matches for individuals who are not present in the target set. For each choice of the match score threshold, we compute precision, recall (true positive rate), and false positive rate as follows.

Let *n*_1_ and *n*_2_ be the number of individuals in Set 1 and Set 2, respectively. For Set 1, let TP_1_ (“true positives”) or FP_1_ (“false positives”) be the number of individuals who are correctly or incorrectly linked, respectively, with a match score above the threshold. For Set 2, let FP_2_ (“false positives”) be the number of individuals who were incorrectly linked with a match score above the threshold. Note that all individuals in Set 1 are technically considered “positives” since there exists a true match in the target set. Individuals represented by FP_1_ are an exception, who are positives that are converted to negatives due to problematic match scores. All individuals in Set 2 are considered “negatives” since a true match does not exist. Given these terms, we calculate

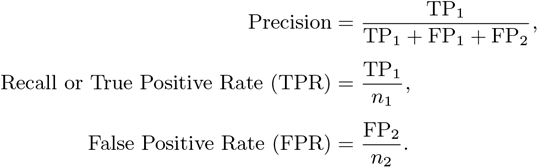

For FPR, we do not consider FP_2_ to keep the denominator of *n*_2_ fixed across different methods.

The receiver-operating-characteristic (ROC) and precision-recall (PR) curves based on the metrics above are reported in Fig. 5. For each split and method, we interpolate PR curves from 0 to the maximum recall on the *x*-axis for that particular split. The ROC curve is similarly interpolated by considering all FPRs from 0 to 0.99.

We note that our modified metrics for linking performance result in ROC and PR curves that do not end at the typical end points, i.e., (1,1) and (1,0), respectively. This is because there are certain individuals (FP_1_) who are incorrectly classified regardless of the decision threshold since the score for the true match is less than the score for the top match for these individuals.

The AUPRC and AUROC metrics reported for each curve considers an *x*-axis threshold *α* up to which the area under the curve (AUC) is computed and then rescaled by 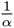. We adopt this truncation approach to equalize the comparisons across methods, given that the location of the end point of each curve (corresponding to a decision threshold that obtains the maximum recall) is different across methods due to the FP_1_ issue describe above. For AUROC, we chose *α* = 0.97 as the FPR threshold and for AUPRC, we chose *α* = 0.85 as the recall threshold, based on the lowest end point of the curves.

## Supplemental Note 3: Forward-backward algorithm for DSM

Recall that during inference, we wish to calculate

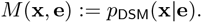

Our haplotype-level approximation factorizes this term as:

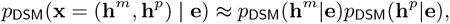

where

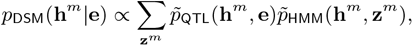

and analogously for *p̃*_DSM_(**h***^p^|***e**). Note that *p̃*_QTL_ represents (unnormalized) probabilistic factors capturing eQTL associations, and *p̃*_HMM_ represents HMM factors—i.e., transition and emission probabilities over the genotypes and hidden states only.

Here we describe a forward-backward algorithm for computing *p̃*_DSM_(**h***^m^|***e**), which is the same as that for **h***^p^*. First, note that

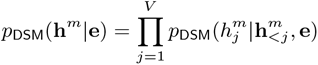

by chain rule, where **h_<j_***^m^* denotes all genotypes that precede the *j*-th position. Each term can be computed using the following relation based on the independence structure of DSM:

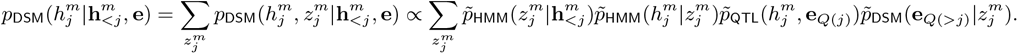

We denote the set of eGenes associated the *j*-th eQTL as **e***_Q_*_(*j*)_ and those associated with any of the subsequent eQTLs as **e***_Q_*_(*>i*)_. Our modification of the standard forward-backward algorithm therefore calculates the “forward” probability, i.e., 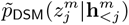, and the “backward” probability, i.e., 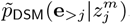, for all j via dynamic programming. The forward term is analogous to the Li-Stephens model, whereas the backward term newly incorporates information from the observed gene expression. Once these terms are computed, the above equation can be used to compute the full conditional likelihood *p*_DSM_(**h***^m^|***e**) as desired.

More detailed computational steps of the forward-backward algorithm are as follows. We begin by defining a dynamic programming matrix *α ∈* [0, 1]*^V^ ^×R^*, where *V* is the number of eQTLs and *R* is the number of reference haplotypes, that stores the (unnormalized) forward probabilities of *z_j_^m^* for each eQTL *j*. Our initial belief over *z*_1_*^m^*, the hidden state for the first eQTL, is uniformly initialized:

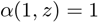

for all *z ∈ {*1*, …, R}*. Then, sequentially for each *j ∈ {*2*, …, V }*, we compute

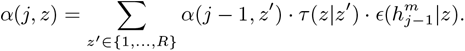

Here, *τ* and *ɛ* indicates the transition and emission probabilities of the HMM component of DSM, respectively. After each step, we rescale *α*(*j, z*) to prevent an underflow.

For backward probabilities, we define a dynamic programming matrix *β ∈* [0, 1]*^V^ ^×R^* that stores probabilities over *z_j_^m^* for each eQTL *j*. Specifically, we begin by setting *β*(*V, z*) = 1 for all *z ∈ {*1*, …, R}*. Then, for all *j ∈ {V −*1*, …,* 1*}*, we set:

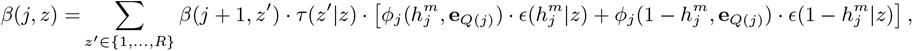

where *ϕ_j_* indicates the eQTL factor at position *j*, which we introduced to model the association between each eQTL and a corresponding set of eGenes.

We also note that while the transition factor *τ* (*z|z^′^*) is näıvely quadratic in the number of reference haplotypes, *τ* (*z|z^′^*) has a structured matrix form:

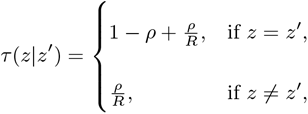

where *ρ* is the recombination rate between the two eQTLs. Thus, the matrix is symmetric and constant everywhere but the diagonal, which enables an efficient multiplication with the transition matrix, linear in the number of reference haplotypes.

The computational cost of the above computation is *O*(*R*) for each of V eQTLs, resulting in the overall complexity of *O*(*V R*). Our memory requirement also has the same complexity, since the sizes of *α* and *β* are both *O*(*V R*). Thus, our runtime and memory grow linearly in both the number of eQTLs (window size) and the number of reference haplotypes. We provide empirical results illustrating the linear memory scaling of our model in Supplemental Fig. S4.

---

**Supplemental Figure S1:**
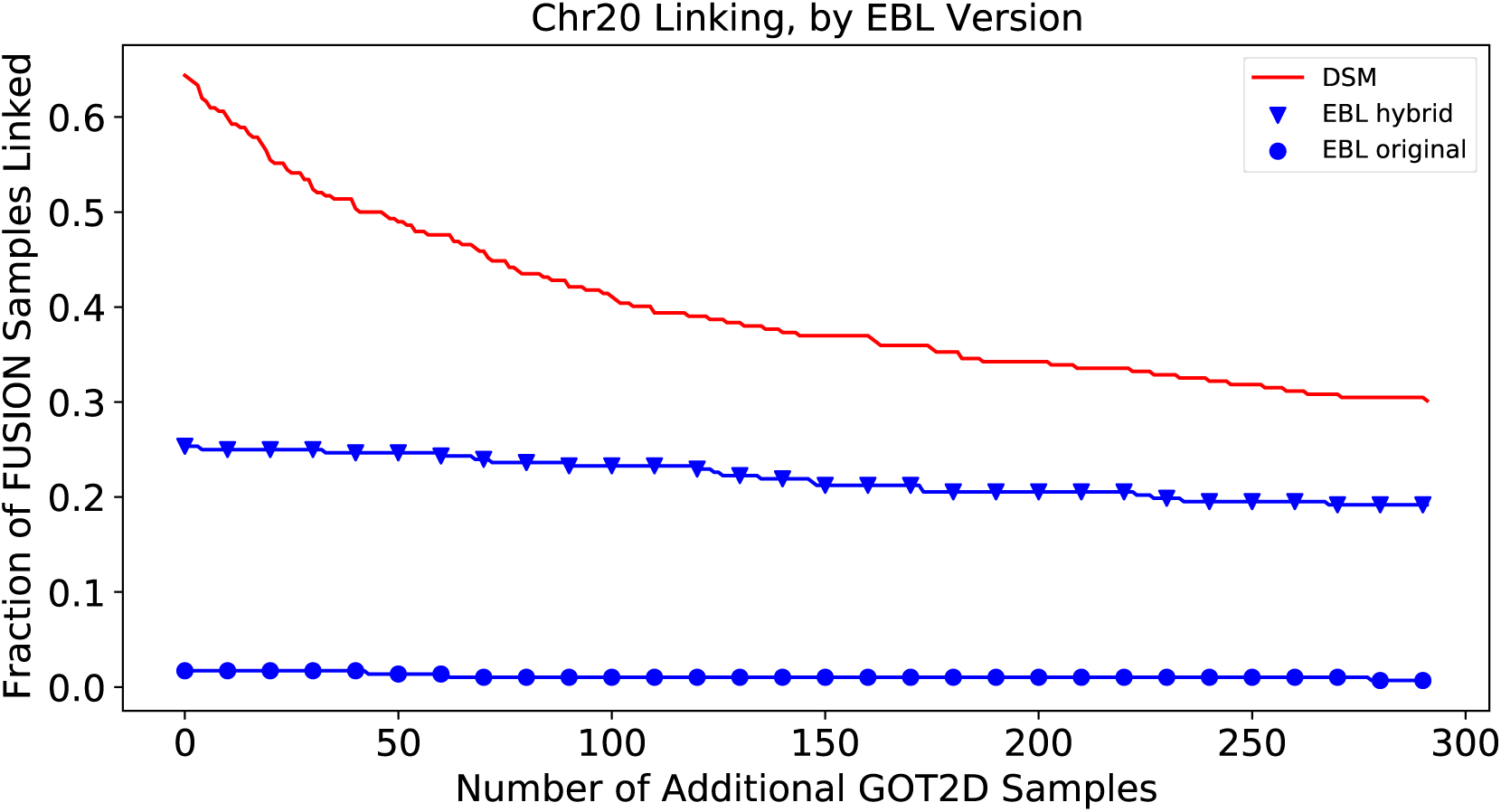
Our hybrid EBL model outperforms the original EBL model. The original EBL models uses an uninformed prior *p*(**x**) estimated from training data MAFs for those individuals and eQTLs where expression values are not extreme (circle). Using a hybrid approach enables a best-of-both-worlds scenario, in which eQTLs not associated to extreme expression values are still given an informative, non-uniform, posterior probability (triangle). The hybrid approach collapses to the GNB model for such eQTLs. Note that the original EBL model uses a sparse set of SNPs across the whole genome, which enables them to achieve high accuracy by using few highly informative SNPs. However, on a single chromosome there are unlikely to be a sufficient number of highly informative SNPs.

**Supplemental Figure S2:**
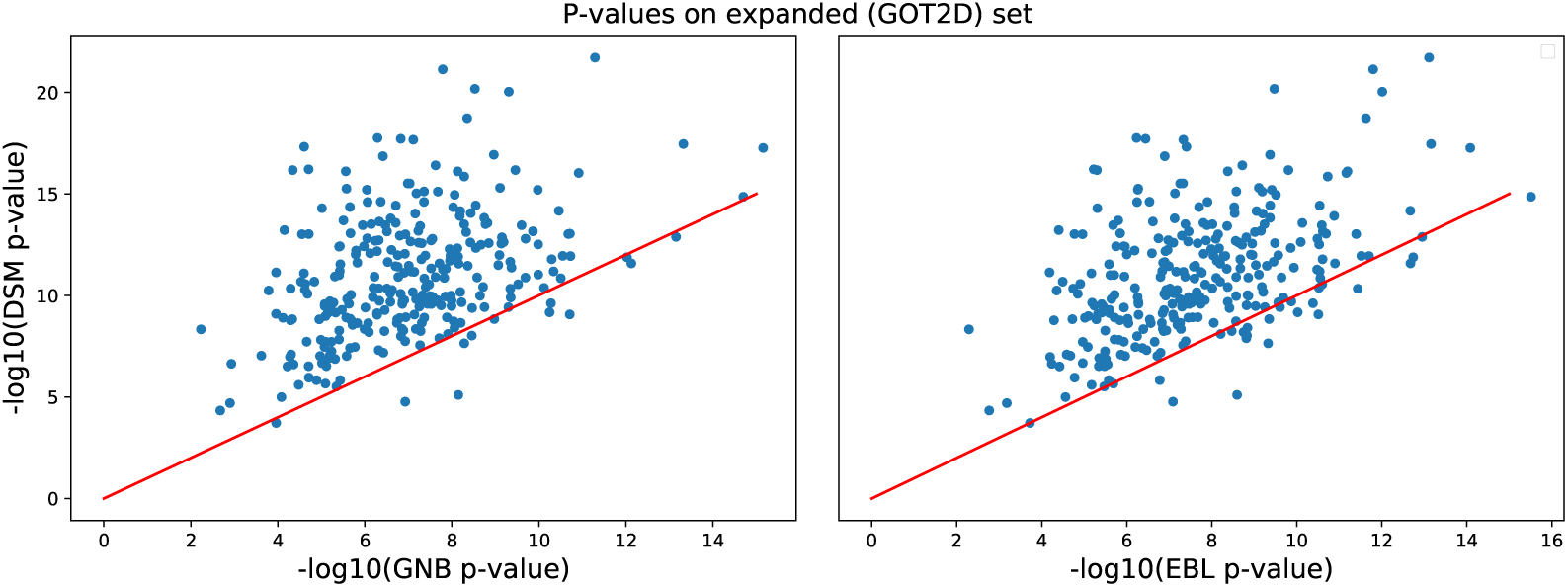
DSM model-based *p*-values are lower than GNB and EBL on expanded GOT2D dataset. When including the expanded set of mismatching GOT2D genotypes for the matching score null distribution, we observe that DSM is able to consistently generate lower *p*-values than both previous approaches for all but a few individuals.

**Supplemental Figure S3:**
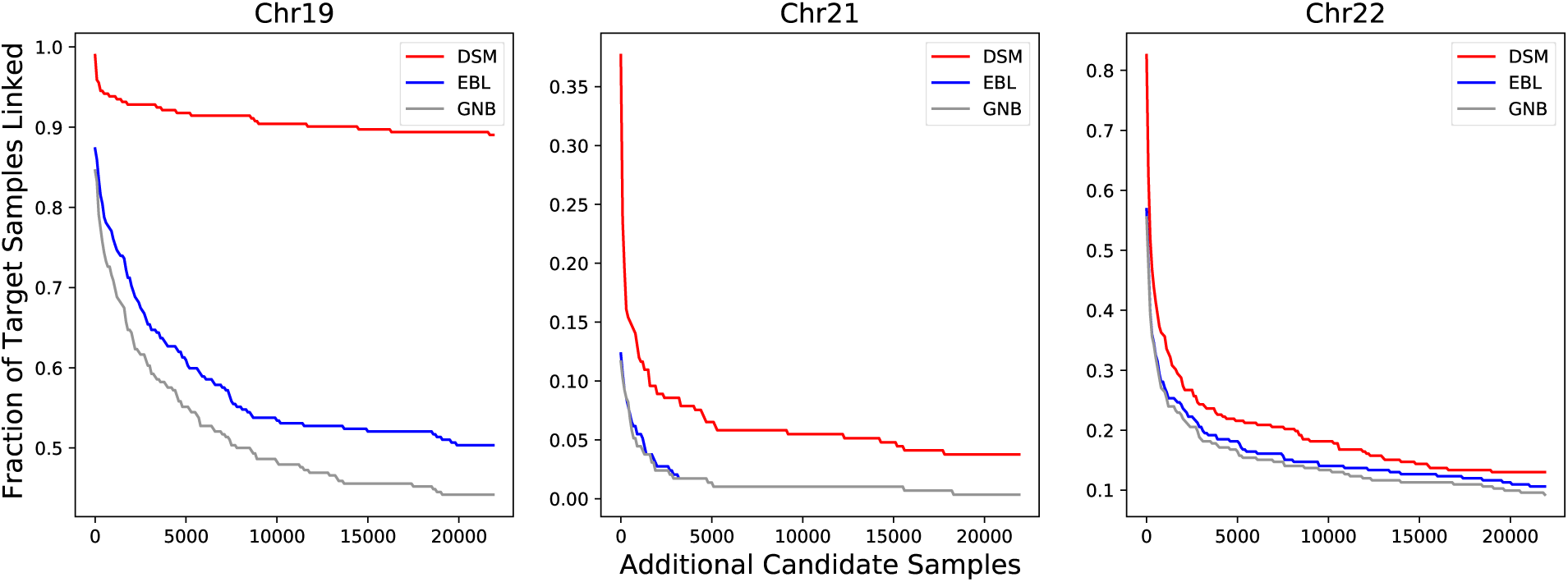
DSM links a greater fraction of individuals on massive candidate genotype sets. On chromosomes 19, 21, and 22, we observed that DSM linked a substantially larger fraction of individuals when including *∼*22k additional genotypes from HRC. This gap is largest in Chromosome 19, where DSM links *>*45% more individuals than EBL and GNB. The results for Chromosome 20 and for all four chomosomes combined are included in Figure 4.

**Supplemental Figure S4:**
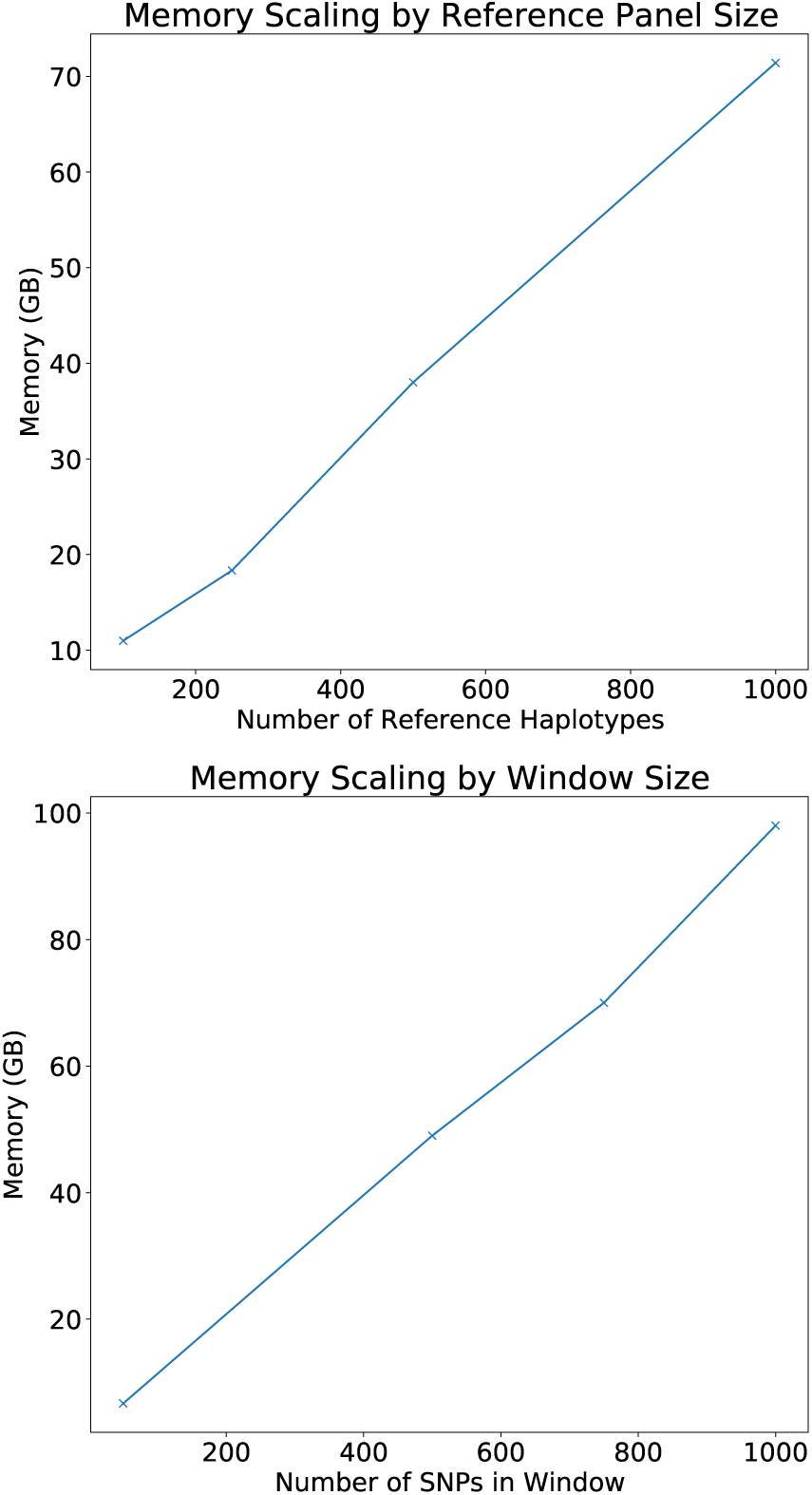
Memory requirement of DSM scales linearly in window and reference panel sizes. Our haplotype-level factor approximation enables us to use a far larger reference panel without quadratic memory requirements (**top**). We also enable linear scaling in memory in the size of windows considered (**bottom**). In practice, we used a reference panel of 1000 haplotypes and window size of 750 eQTLs, which allowed us to train the DSM with a practical memory requirement of *∼*75GB. These are notably larger numbers than those required for GNB and EBL, which can each be done on the order of minutes for a full chromosome with less than 5GB memory. However, note that it is reasonable to consider an adversary with sufficient computational resources for the purpose of assessing the risk of linking attacks.

**Supplemental Figure S5:**
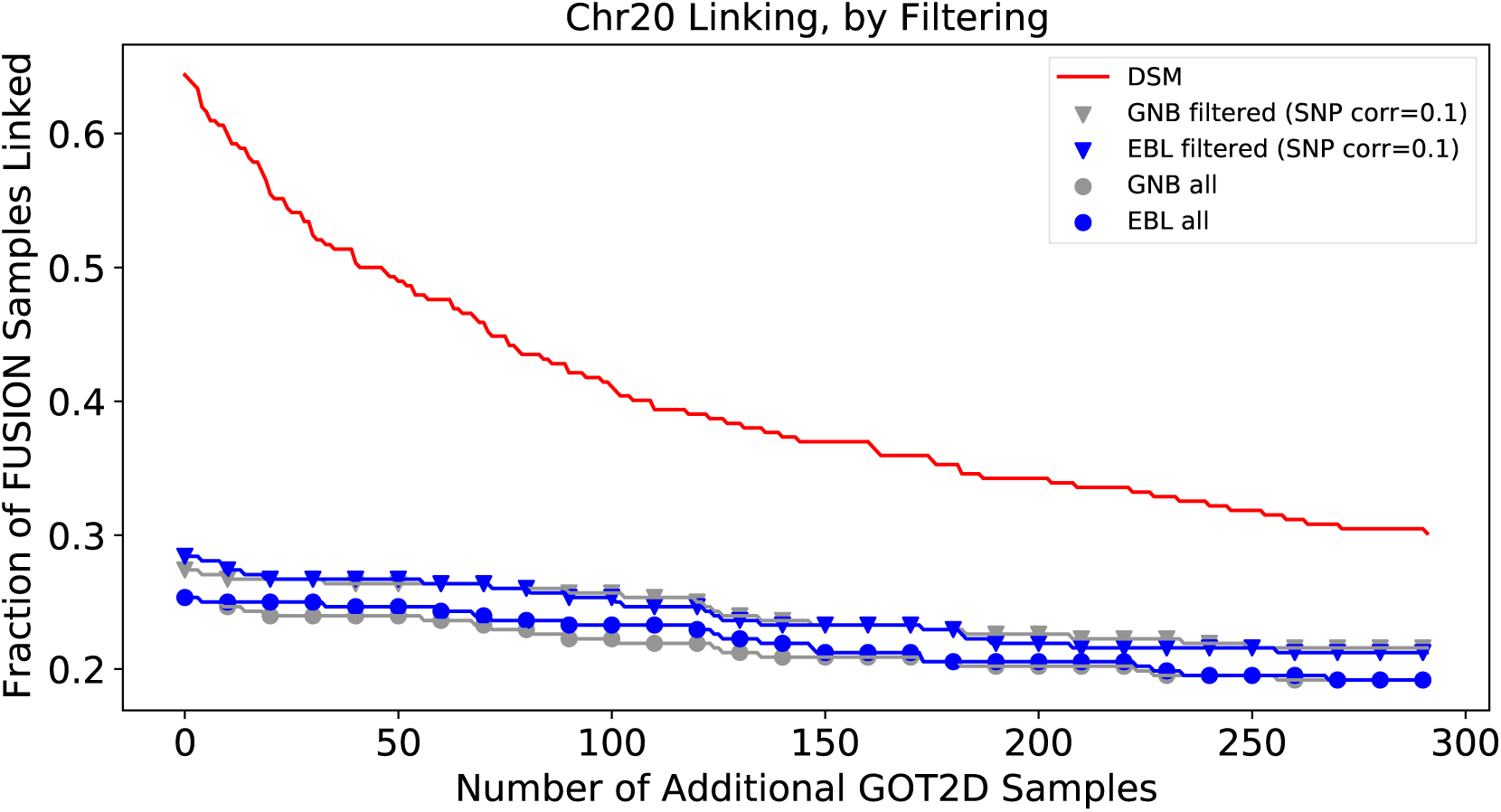
Filtering to remove correlated eQTLs helps GNB and EBL models. EBL and GNB both assume some level of pairwise independence between eQTLs, as including all eQTLs can lead to miscalibrated matching score probabilities. For both these models, the linking accuracy pre-filtering (circle) is worse than the linking accuracy post-filtering (triangle). The SNP set is greedily pruned until no two SNPs have pairwise correlation greater than 0.1.

**Supplemental Figure S6:**
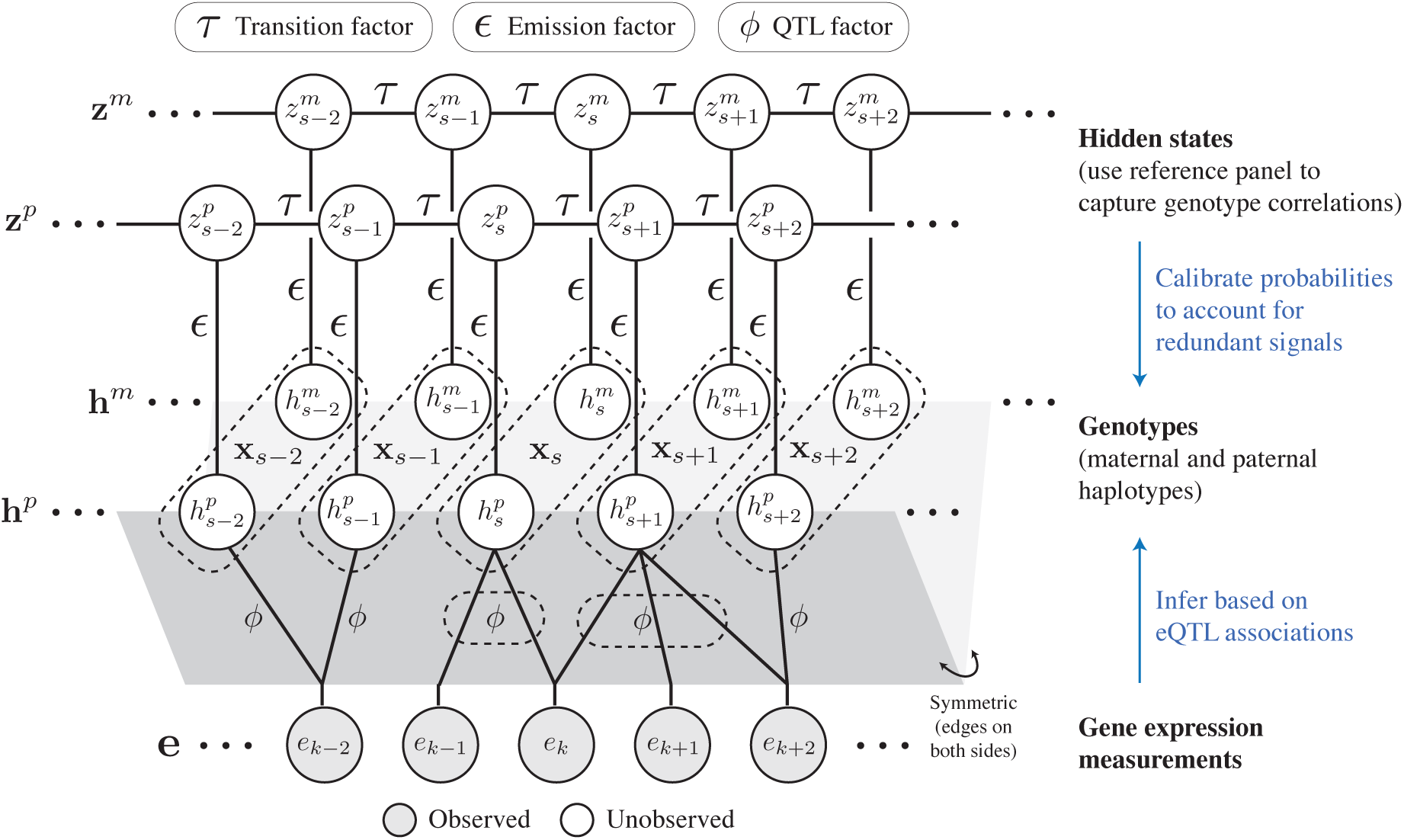
Graphical representation of DSM. Solid edges indicate factors in the conditional distribution *p*(**x***|***e**), including transition, emission, and QTL factors. **h** corresponds to the haplotype, of which there are two (together denoted **x**, indicated by dotted lines), while **e** corresponds to the expression levels of individual genes. The *ϕ* factors, some of which involve multiple genes (dotted lines), relate predictive signals in the gene expression data to the genotypes (pairs of alleles). To enable the use of large reference panels, we predict the haplotype allele and copy this prediction twice (gray foldout). The *τ* and *ɛ* factors respectively correspond to the recombination and emission probabilities in the standard Li-Stephens model. **z***^m^* and **z***^p^* index into the reference panel of haplotypes, whose posteriors, conditioned on observed gene expression values, are updated via the forward-backward algorithm.

**Supplemental Table 1:** DSM’s reverse linking is more accurate than linking based on predicted gene expression. We report DSM’s accuracy in linking genotype profiles to matching expression profiles (i.e., in the reverse direction from our primary evaluation setting), compared to an alternative approach based on a state-of-the-art gene expression prediction method, MetaXcan [Barbeira et al., *Nature Communications*, 2018]. The evaluations are performed on the FUSION dataset. We used the pretrained MetaXcan model on GTEx v8 muscle-skeletal tissue dataset, the same dataset used to train DSM. Both Pearson and Spearman’s correlation coefficients between the predicted and observed expression profiles are considered as the match score for MetaXcan. DSM leads to more accurate linking than matching based on the predicted gene expression levels from MetaXcan.

| Chromosome | MetaXcan (Pearson) | MetaXcan (Spearman's) | DSM |
| --- | --- | --- | --- |
| 19 | 114 (39.0%) | 51 (17.5%) | **291** (99.7%) |
| 20 | 15 (5.1%) | 6 (2.0%) | **240** (82.2%) |
| 21 | 3 (1.0%) | 4 (1.4%) | **157** (53.8%) |
| 22 | 24 (8.2%) | 13 (4.5%) | **263** (90.0%) |
| All 4 combined | 171 (58.6%) | 97 (33.2%) | **291/292** (99.7%) |

